# A CRISPR/Cas9-generated mutation in the zebrafish orthologue of *PPP2R3B* causes idiopathic scoliosis

**DOI:** 10.1101/413526

**Authors:** Marian Seda, Berta Crespo, Michelangelo Corcelli, Daniel P. Osborn, Dagan Jenkins

## Abstract

Idiopathic scoliosis (IS) is the deformation and/or abnormal curvature of the spine that develops progressively after birth. It is a very common condition, affecting approximately 4% of the general population, yet the genetic and mechanistic causes of IS are poorly understood. Here, we focus on *PPP2R3B*, which encodes a protein phosphatase 2A regulatory subunit that has been linked to regulation of autophagy. We found that *PPP2R3B* is expressed at sites of chondrogenesis within human foetuses, including the vertebrae. We also demonstrated prominent expression in myotome and muscle fibres in human foetuses, and zebrafish embryos and adolescents. As there is no rodent orthologue of *PPP2R3B*, we used CRIPSR/Cas9-mediated gene-editing to generate a series of frameshift mutations in zebrafish *ppp2r3b*. Adolescent zebrafish that were homozygous for this mutation exhibited a fully penetrant kyphoscoliosis phenotype which became progressively worse over time, mirroring IS in humans. These defects were associated with reduced mineralisation of vertebrae, resembling osteoporosis. Electron microscopy demonstrated abnormal mitochondria adjacent to muscle fibres, suggestive of abnormal mitophagy.

## Introduction

Scoliosis is the lateral deformation and curvature of the spine which affects approximately 4% of the general population (Cheng et al. 2015). This can result from a primary defect of the bone that constitutes the vertebrae, and indeed some mouse models of scoliosis exhibit alterations of osteoblasts and chondrogenesis (Liang et al. 2018). Scoliosis can also arise secondarily to defects in proprioception (Blecher et al. 2017), which refers to the body’s sense of position and reaction to external stimuli. This is driven by connections between the interneurons that relay signals such as pain to motor neurons within the spinal cord, which in turn control muscle activity. Defects in these neuronal connections and activities can lead to progressive idiopathic scoliosis (IS). The association of muscular dystrophy with IS also emphasises the importance of musculature in maintaining spinal integrity. Another mechanism thought to lead to IS is abnormal cerebrospinal fluid flow (Grimes et al. 2016). It is therefore important to define the primary cellular origins of different genetic forms of scoliosis.

*PPP2R3B* which encodes the PR70 protein, a regulatory subunit of the heterotrimeric protein phosphatase 2A holoenzyme. Relatively little is known about PR70 function, although it was identified as interacting with the origin of replication complex component, CDC6, and has also been linked to regulation of autophagy (Yan et al. 2000; Dovega et al. 2014; Pengo et al. 2017). PR70 has also been proposed to act as a tumour suppressor by regulating firing of DNA replication origins (Van Kempen 2016). In this study we have used CRISPR/Cas9-mediated gene-editing to create a frameshift mutation in the zebrafish *ppp2r3b* gene. Homozygous mutant larvae appeared normal, while adolescents developed progressive kyphoscoliosis reminiscent of IS. *PPP2R3B* showed prominent expression in axial muscle in human foetuses, zebrafish embryos and adolescents. The mutant kyphoscoliosis phenotype is associated with reduced vertebral bone mineralisation and a muscular dystrophy-like phenotype. These data identify *PPP2R3B* as a new molecular target in the pathogenesis of human IS.

## Results

### Expression of *PPP2R3B/ppp2r3b* in human foetuses and zebrafish

To begin to investigate the requirement for PR70 in vertebrates, we looked for expression of *PPP2R3B* in human foetuses. Using *in situ* hybridisation, we noted some locations of *PPP2R3B* expression that are potentially relevant to scoliosis. This included interneuron and motor neuron precursors within the neural tube, dorsal root ganglia and myotome (Figure 1A). As a control, we used a GFP antisense probe with the same length and GC content as the *PPP2R3B* probe, which gave no signal (Figure 1B). Expression was also noted within the vertebrae as well as Meckel’s cartilage (Figure 1C,E). *PPP2R3B* transcripts were detected within cartilage condensations suggestive of a role in chondrogenesis, although it was not expressed within the perichondrium where osteoblast precursors reside prior to their migration into the cartilage matrix (Figure 1C’,E’). In both of these locations, *PPP2R3B* expression colocalised with *SOX9* on adjacent sections (Figure 1D’,G’). Within Meckel’s cartilage, chondrocytes within cartilage condensations also expressed *SOX10*, as did the perichondrium (Figure 1F’). *SOX10* is a marker of neural crest stem cells confirming the contribution of this lineage to skeletal elements within the jaw. We note that *PPP2R3B* expression was quite broad, albeit with accentuation of signal in certain locations, including myotome and vertebral chondrocytes, as shown by intermediate power images (Figure 1A’,C”).

**Figure 1.**
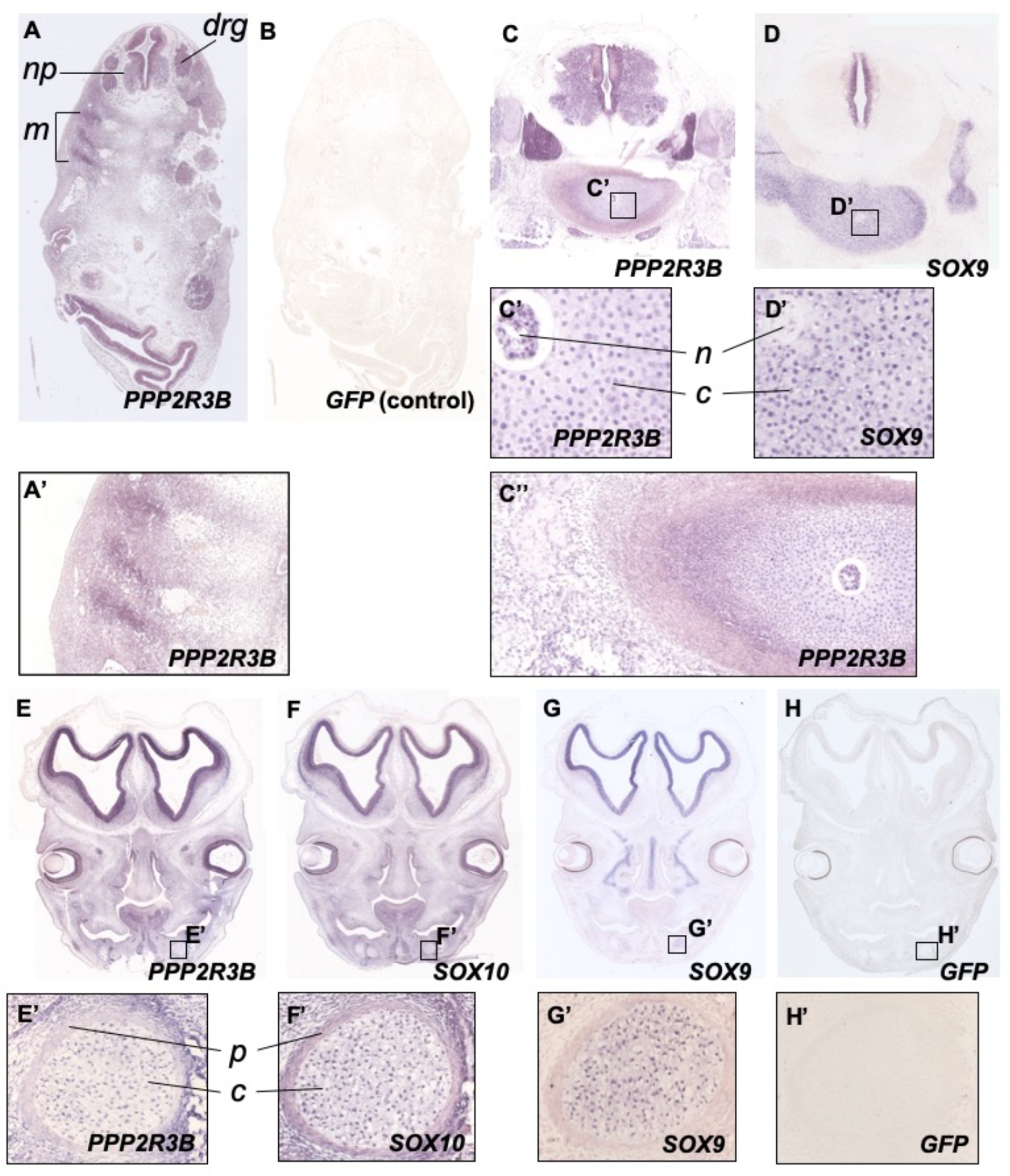
*PPP2R3B* is expressed at sites of chondrogenesis in normal human foetuses. Expression of *PPP2R3B, SOX9* and *SOX10* in normal human foetuses at Carnegie stages (CS) 17 (**A,B**) 23 (**C,D**) and 22 (**E-H**). (**A,B**) *In situ* hybridisation at low power showing expression of *PPP2R3B* within interneuron/motor neuron precursors (*np*), dorsal root ganglia (*drg*) and myotome (*m*) but no signal generated using a GFP negative control. (**C,D**) Expression of *PPP2R3B* in vertebral bodies. Insets show regions magnified in **C’** and **D’**. Note expression in chondrocytes (*c*). *PPP2R3B* is also expressed in the notochord (*n*) whereas *SOX9* is not. (**E-H**) Expression of *PPP2R3B* within Meckel’s cartilage (insets magnified in **E’-H’**). Note expression co-localises with *SOX10* and *SOX9* within chondrocytes (*c*) but expression is absent from perichondrium (*p*).

By aligning the human PR70 protein sequence to the zebrafish translated genome, we identified only a single orthologue with significant similarity, and the genomic locus encoding Ppp2r3b showed conserved synteny with their mammalian counterparts (Figure 2A). Orthologues of neither *PPP2R3B* nor the adjacent gene, *SHOX*, are found in rodents. We therefore analysed expression of the orthologous zebrafish *ppp2r3b* gene by *in situ* hybridisation (Figure 2B,C). At 24 hours post-fertilisation (hpf) we noted repeated chevron-shaped patterns of expression along the trunk of the embryo representing the somites which will go on to form muscle, ribs and vertebrae. Within the craniofacial region, we also noted expression throughout the branchial arches and somites at 24 hpf, in the vicinity of actively migrating NCSCs (Figure 2D).

**Figure 2.**
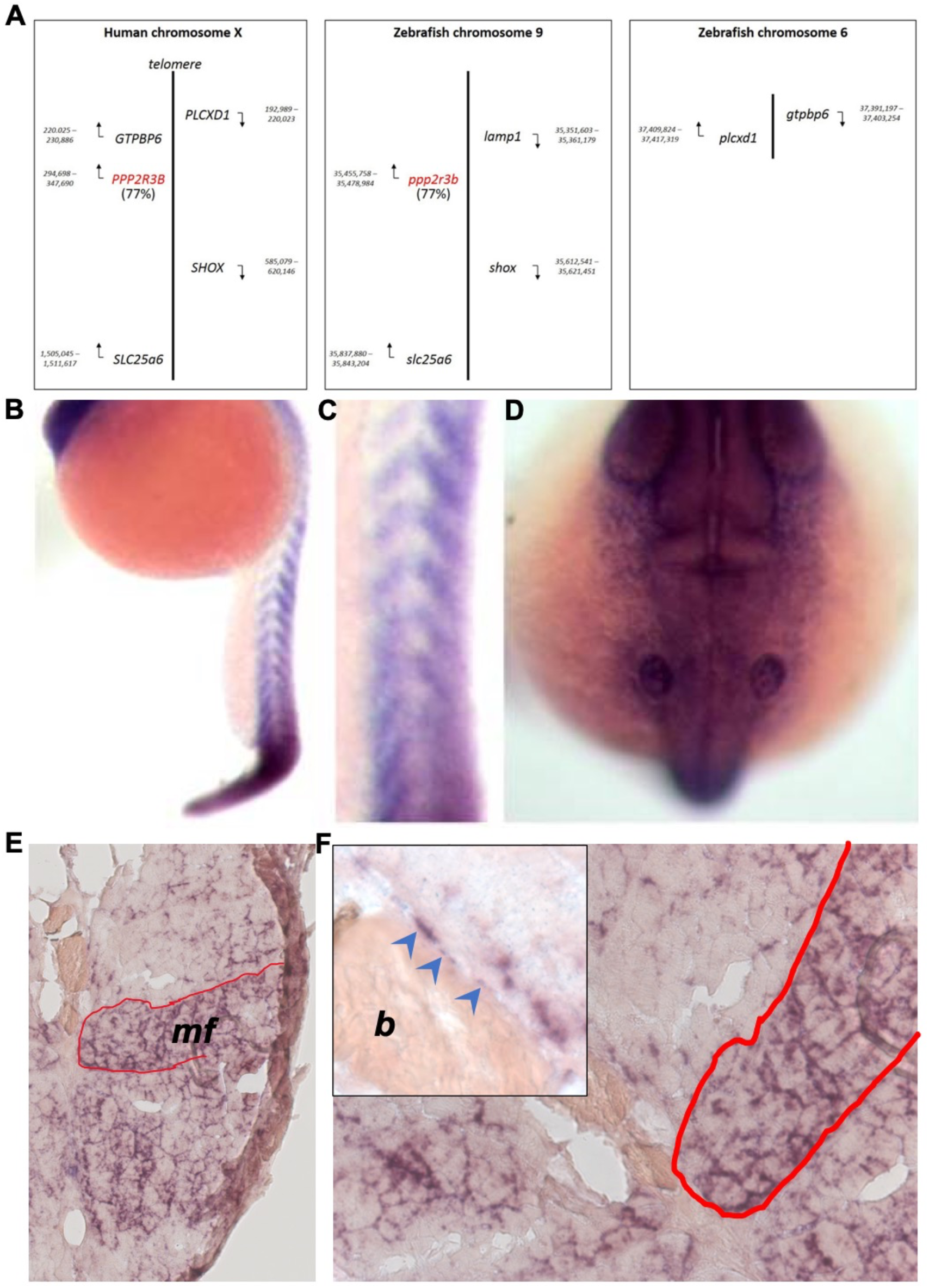
Expression of *ppp2r3b* in zebrafish. **(A)** Schematic showing conservation of synteny between zebrafish and human *ppp2r3b/PPP2R3B*. On the left is part of the pseudoautosomal region on the human X-chromosome giving the co-ordinates of each gene, and indicating the strand from which is gene is transcribed (arrows). The percentage amino acid conservation between human and zebrafish PPP2R3B/Ppp2r3b is indicated. *Middle-* zebrafish chromosome 9 showing conserved synteny for *slc25a6, shox* and *ppp2r3b*, as well as the adjacent *lamp1* gene. *Right-* zebrafish chromosome 6 showing limited conserved synteny of *plcxd1* and *gtpbp6*. (**B-D**) *In situ* hybridisation showing expression of *ppp2r3b* at 24 hpf. High-power view showing expression in somites (**C**) and neuroectoderm/migrating neural crest (**D**). (**E,F**) Transverse section showing widespread expression of *ppp2r3b* at 36 dpf throughout axial muscle fibres (*mf*). Mineralised bone (*b*) within vertebral centra showing expression of *ppp2r3b* in squamous chordoblast cells (arrowheads).

We also analysed expression in zebrafish at 36 dpf of age. We noted prominent expression throughout the axial muscle (Figure 2E,F). In contrast there was only limited expression in proximity to mineralised bone - we did identify expression in squamous chordoblast (osteoblast)-like cells which are present on the vertebral bone surface, as previously reported (Bensimon-Britto et al.), although it is noteworthy that these cells were very rare (Figure 2F, inset).

### Generation of *ppp2r3b* mutant zebrafish using CRISPR/Cas9 gene-editing

To generate a genetic model of *ppp2r3b* loss-of-function, we used CRISPR/Cas9 gene-editing to target this gene using a sgRNA located within exon 2 (Figure 3A). This sgRNA was located on the forward strand and had no self-complementarity or predicted off-target sites according to the chopchop tool (http://chopchop.cbu.uib.no/). An *Mse I* restriction site was located within the sgRNA binding site which allowed us to monitor the efficiency with which indels were introduced at this location. Co-injection of Cas9 RNA and a sgRNA targeting *ppp2r3b* resulted in mosaic mutations (data not shown). Direct sequencing of a selection of cloned mutations from mosaic F0 embryos at 24 hpf was consistent with many previous reports in zebrafish, showing that CRISPR/Cas9 typically produces complex indels involving deletions of between 2-18 nucleotides (Figure 3A).

**Figure 3.**
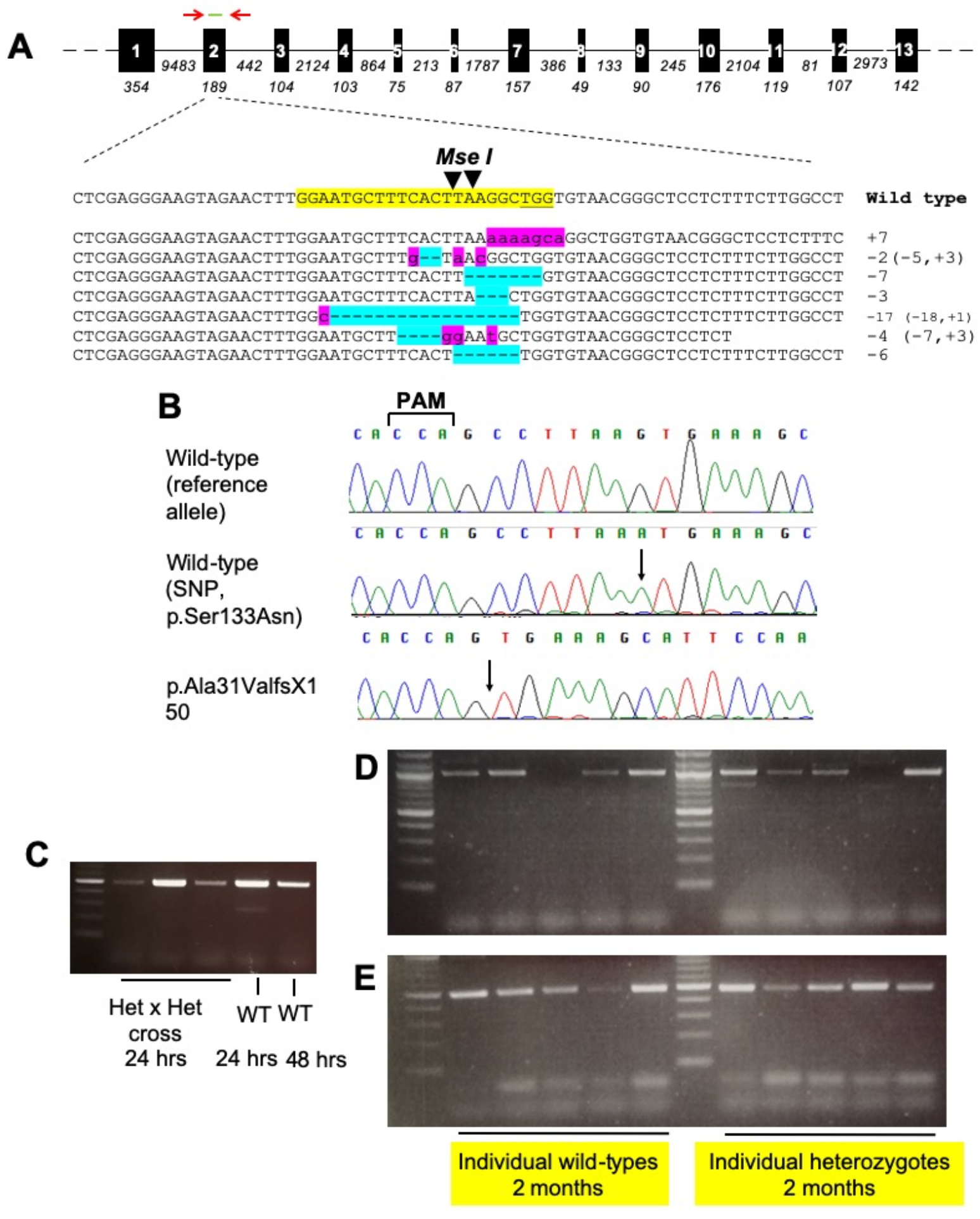
Targeting *ppp2r3b* in zebrafish using gene-editing. (**A**) *ppp2r3b* gene structure showing the location of primers used for genotyping (red arrows) and sgRNA (green) used for gene-editing. Below, examples of mutant sequence reads cloned from pooled F0 embryos. The sgRNA site is highlighted in yellow within the wild-type sequence. Inserted and deleted nucleotides are highlighted in pink and blue, respectively. (**B**) Sequence chromatograms showing the homozygous wild-type reference and alternative alleles, and the homozygous and heterozygous mutant reads. (**C,D**) RT-PCR showing expression of *ppp2r3b* using primers located within exons 1-3 (**C,E**) or exons 1-7 (**D**) at the indicated stages.

We have now outcrossed these F0 mosaics and their progeny for more than 5 generations to achieve germline transmission and to avoid possible off-target mutations. We isolated a line of zebrafish carrying a 7 bp deletion in exon 2 of *ppp2r3b* which results in the frameshift mutation p.Ala31ValfsX150 (Figure 3B). Homozygous mutants are hereafter referred to as *ppp2r3b^−/−^*. During the course of our breeding and genotyping, we noted a single nucleotide polymorphism (SNP) located within the sgRNA binding site which is not present on publicly available databases. This SNP encodes the single amino acid substitution p.Ser33Asn. In all subsequent analyses, we selected only heterozygotes whose wild-type allele encoded the reference SNP at this location in our breeding population. RT-PCR using primers located in exons 1-3 or 1-7 confirmed that all *ppp2r3b* transcripts generated from pooled 24 hpf embryos from a hetxhet incross or individual homozygous mutant animals at 48 dpf of age were of the predicted size, as in wild-types, and qRT-PCR demonstrated that transcript levels in mutants were not different from wild-type (Figure 3C-E, Table 1). We noted that there was no deviation from expected Mendelian ratios showing that this mutation does not affect viability (Figure 4). We also endeavoured to generate a pool of homozygous mutant adults with which to breed maternal-zygotic mutant zebrafish. This was not possible, because homozygotes never produced any eggs, and thus we conclude that they are infertile.

**Figure 4.**
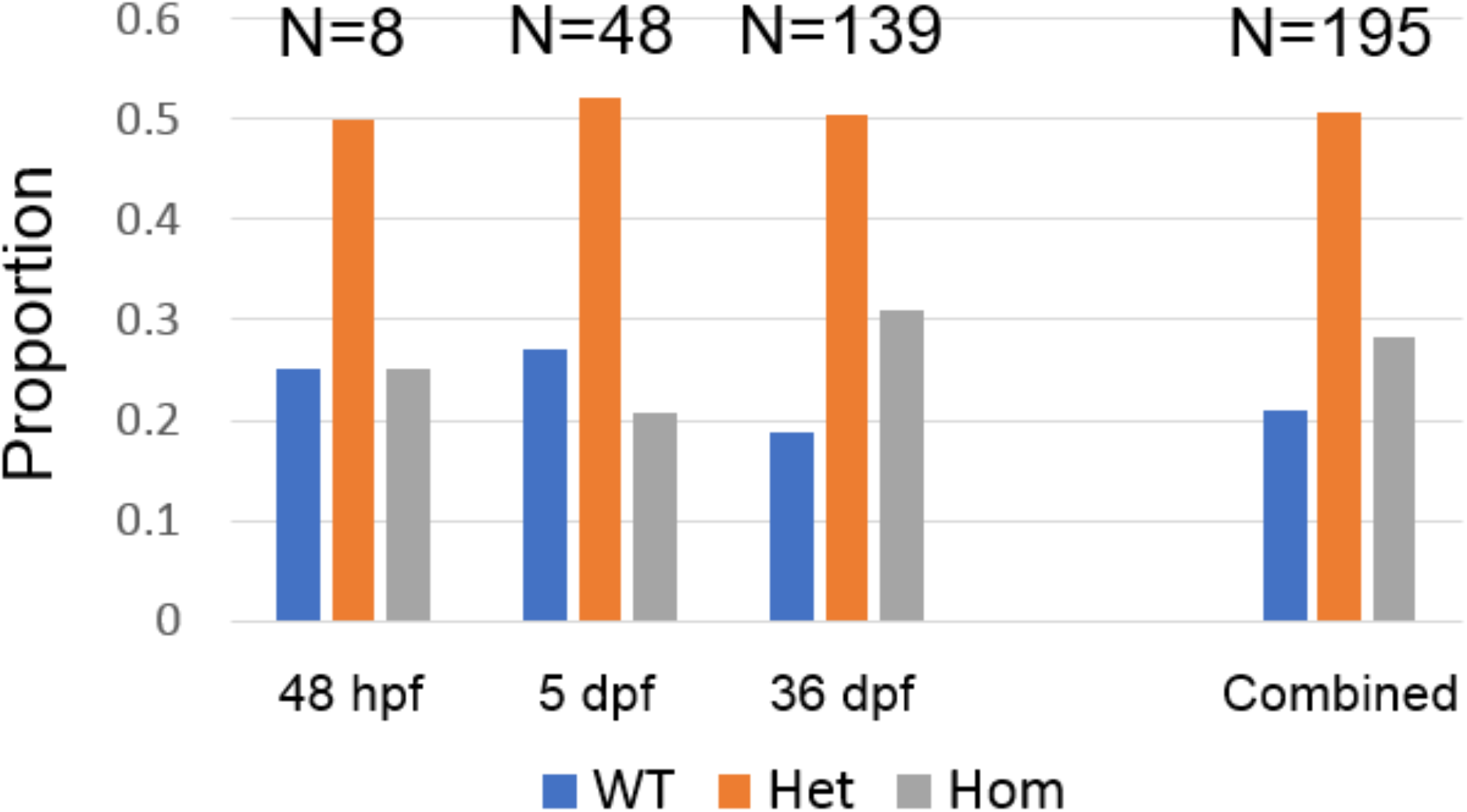
*ppp2r3b* mutants are viable at all ages. Proportions of wild-type (WT), heterozygous (Het) and homozygous (Hom) mutant (*ppp2r3b* ^Ala31ValfsX150^) animals at the stated ages. Total numbers of embryos analysed are indicated.

**Table 1.** qPCR analysis of bone and muscle marker genes.

| Gene | Genotype | Rel. expression | SEM | p-value |
| --- | --- | --- | --- | --- |
| ***Target gene*** |  |  |  |  |
| ppp2r3b (primer set 1) | WT | 1 | 0.20566 | 0.7752 |
| ppp2r3b (primer set 1) | ppp2r3b-/- | 0.91618 | 0.19554 |  |
| ppp2r3b (primer set 2) | WT | 1 | 0.20067 | 0.241 |
| ppp2r3b (primer set 2) | ppp2r3b-/- | 0.68974 | 0.14055 |  |
| ***Muscle marker genes*** |  |  |  |  |
| mstnb | WT | 1 | 0.21493 | 0.0512 |
| mstnb | ppp2r3b-/- | 0.42757 | 0.12753 |  |
| mylz2 | WT | 1 | 0.20609 | 0.1026 |
| mylz2 | ppp2r3b-/- | 0.56843 | 0.11124 |  |
| smyhc1 | WT | 1 | 0.21205 | 0.782 |
| smyhc1 | ppp2r3b-/- | 1.29499 | 1.00891 |  |
| stnnc | WT | 1 | 0.34582 | 0.5894 |
| stnnc | ppp2r3b-/- | 1.93724 | 1.63087 |  |
| myhz2 | WT | 1 | 0.24932 | 0.8358 |
| myhz2 | ppp2r3b-/- | 1.0712 | 0.22007 |  |
| tnni2 | WT | 1 | 0.28223 | 0.6054 |
| tnni2 | ppp2r3b-/- | 0.8193 | 0.18237 |  |
| myog | WT | 1 | 0.2499 | 0.6072 |
| myog | ppp2r3b-/- | 1.18719 | 0.2449 |  |
| tnnc1b | WT | 1 | 0.25356 | 0.2269 |
| tnnc1b | ppp2r3b-/- | 0.64582 | 0.09455 |  |
| ***Bone marker genes*** |  |  |  |  |
| ctsk | WT | 1 | 0.17927 | 0.16 |
|  | ppp2r3b-/- | 1.81554 | 0.49503 |  |
| rank l | WT | 1 | 0.2353 | 0.6865 |
|  | ppp2r3b-/- | 1.15204 | 0.27555 |  |
| col2a1a | WT | 1 | 0.19301 | 0.2527 |
|  | ppp2r3b-/- | 1.61816 | 0.4685 |  |
| mmp2 | WT | 1 | 0.2274 | 0.7332 |
|  | ppp2r3b-/- | 0.89729 | 0.18151 |  |
| mmp9 | WT | 1 | 0.27651 | 0.2306 |
|  | ppp2r3b-/- | 0.61384 | 0.11011 |  |
| mmp13 | WT | 1 | 0.3806 | 0.5372 |
|  | ppp2r3b-/- | 1.36542 | 0.42002 |  |
| timp2a | WT | 1 | 0.19044 | 0.1307 |
|  | ppp2r3b-/- | 0.60948 | 0.13231 |  |
| spp1 | WT | 1 | 0.17135 | 0.6068 |
|  | ppp2r3b-/- | 1.1612 | 0.24748 |  |
| runx2b | WT | 1 | 0.17498 | 0.8269 |
|  | ppp2r3b-/- | 1.05905 | 0.19411 |  |
| sox9b | WT | 1 | 0.15355 | 0.5506 |
|  | ppp2r3b-/- | 1.16335 | 0.21253 |  |

### *ppp2r3b* homozygous mutant zebrafish exhibit a fully penetrant scoliosis phenotype

We did not identify any phenotypic abnormalities in heterozygous or homozygous mutants at larval stages. By 48 dpf, we noted that homozygotes developed severe kyphoscoliosis (Figure 5A,B). The pattern of kyphoscoliosis was very stereotypical, characterised by two ventral curves located within the precaudal and caudal vertebrae at numbers 7-9 and 25, respectively. These ventral curves flanked a dorsal curve located at approximately caudal vertebrae number 18. There was often also a sharp lateral bend within the caudal fin, although this was not as consistent. At this age, wild-type siblings never exhibited kyphoscoliosis and the spine exhibited a very gentle ventral curvature within the precaudal region only. Scoliosis is a common phenotype in old zebrafish, presenting in excess of 18 months of age in our aquatics facility but never earlier than this. This is often associated with *Mycobacterium chelonae* infection, however, ongoing microbiological testing confirms that this species is absent from our facility.

**Figure 5.**
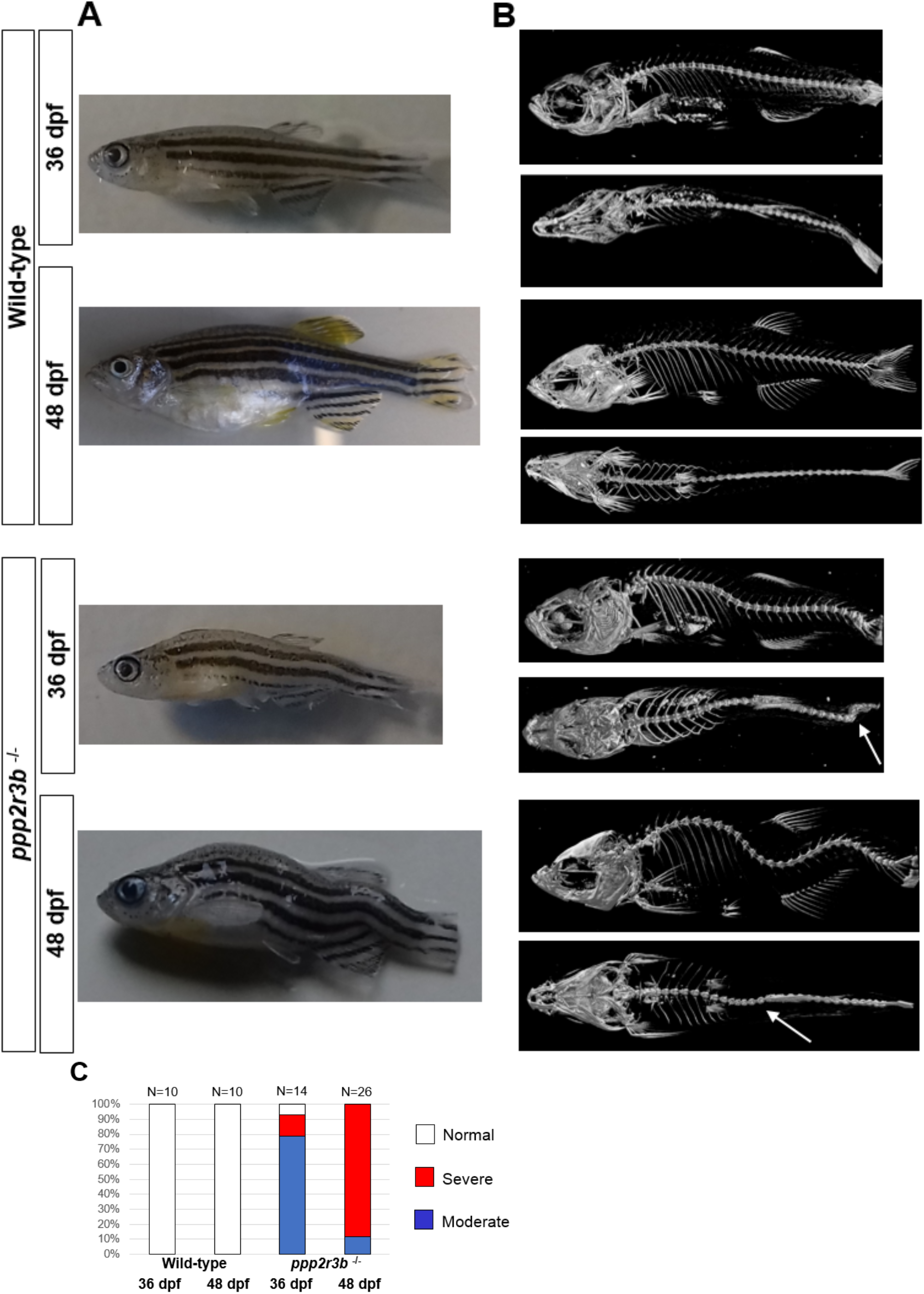
(**A**) Bright field images and (**B**) microCT scans of representative wild-type or homozygous *ppp2r3b^−/−^* mutant zebrafish at 36 or 48 dpf. White arrows in **B** point to sharp lateral curvatures of the spine. (**C**) Quantification of the proportion of animals with mild, moderate or severe kyphoscoliosis.

We monitored the onset and progression of scoliosis in *ppp2r3b^−/−^* mutants. Scoliosis was first seen at 36 dpf (Figure 5; Figure 6A). At this age, the typical presentation was moderate ventral curvature within the precaudal region, with relatively little curvature of the caudal vertebral regions. However, by 48 dpf, the final pattern consisting of two ventral curves and one dorsal curve was apparent. Quantification of the proportion of animals with moderate or severe kyphoscoliosis confirmed that this phenotype became worse with time (Figure 5C). At 48 dpf, the kyphoscoliosis phenotype was present in all homozygous mutants, but not in any wild-type or heterozygous siblings. Therefore, *ppp2r3b^−/−^* mutant zebrafish exhibit adolescent onset and progressive kyphoscoliosis, which is fully penetrant and reminiscent of human IS.

**Figure 6.**
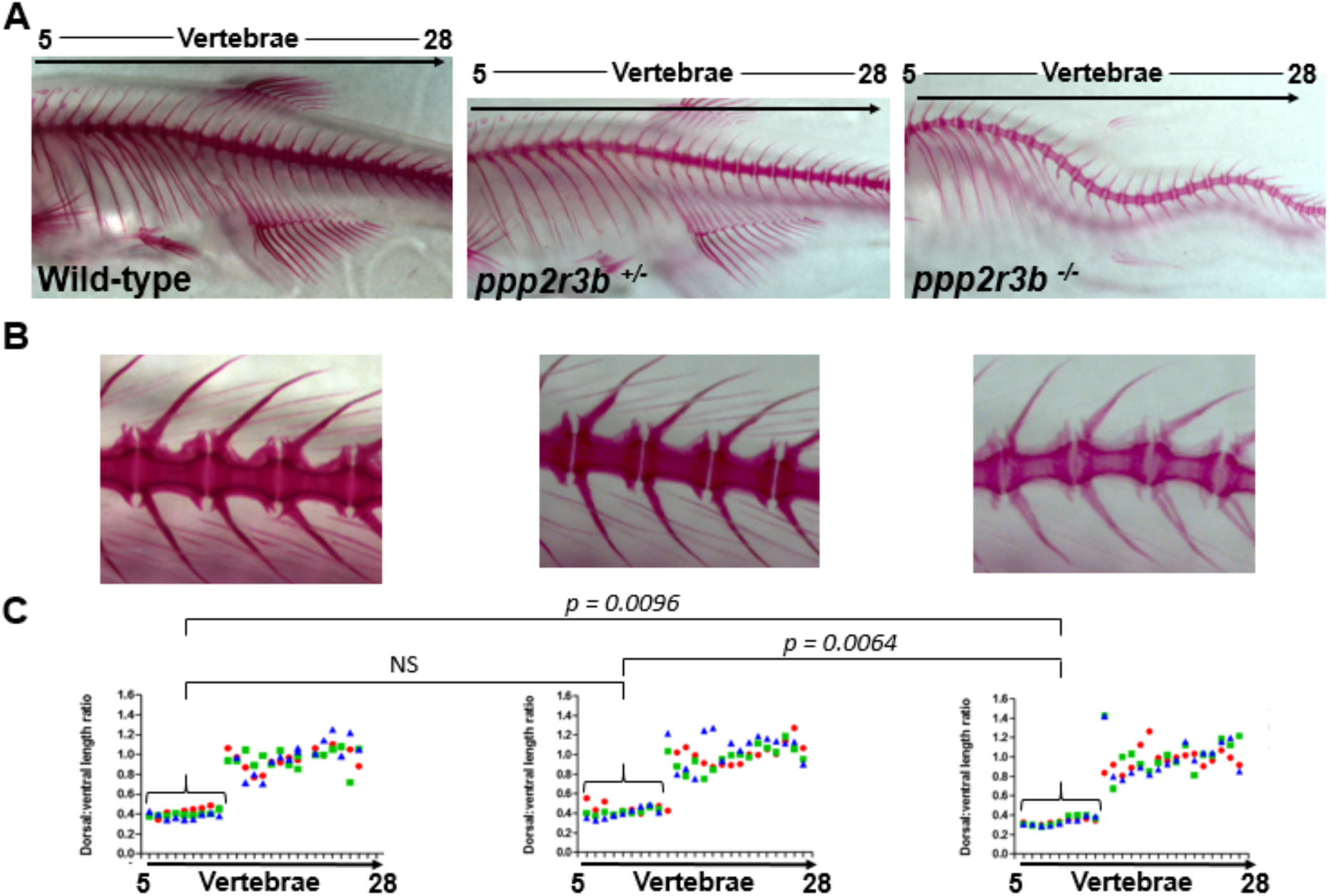
Morphometric analysis of mineralised bone. (**A,B**) Alizarin red staining of vertebrae 36 dpf zebrafish of the indicated genotypes. (**C**) Quantification of the neural spine:hemal spine/rib length ratios for vertebrae 5-28. Data-points for individual animals are indicated by different colours. Mean±standarad deviation of values of these ratios averaged across all ribs for each animal, indicated by the brackets, and subsequently averaged over three animals are 0.401±0.024, 0.423±0.029 and 0.335±0.005 for wild-type, heterozygous and homozygous mutant animals, respectively. This represents a statistically significant difference in homozygotes versus each of the other two genotypes (*t-test*). Tail lengths taking into account spinal curvature were 12.694, 12.872 and 12.749 mm for the representative wild-type, *ppp2r3b+/−* and *ppp2r3b−/−* animals shown, respectively.

To confirm that this phenotype is not the product of off-target mutations we also generated a second frameshift mutant line following microinjection of a ribonucleoprotein complex consisting of sgRNAs targeting exon 1 in complex with Cas9. This generated a 19bp deletion, causing an out-of-frame p.L82fsX24 mutation. Notably, homozygous zebrafish for this mutation also exhibited profound scoliosis (Figure 7).

**Figure 7.**
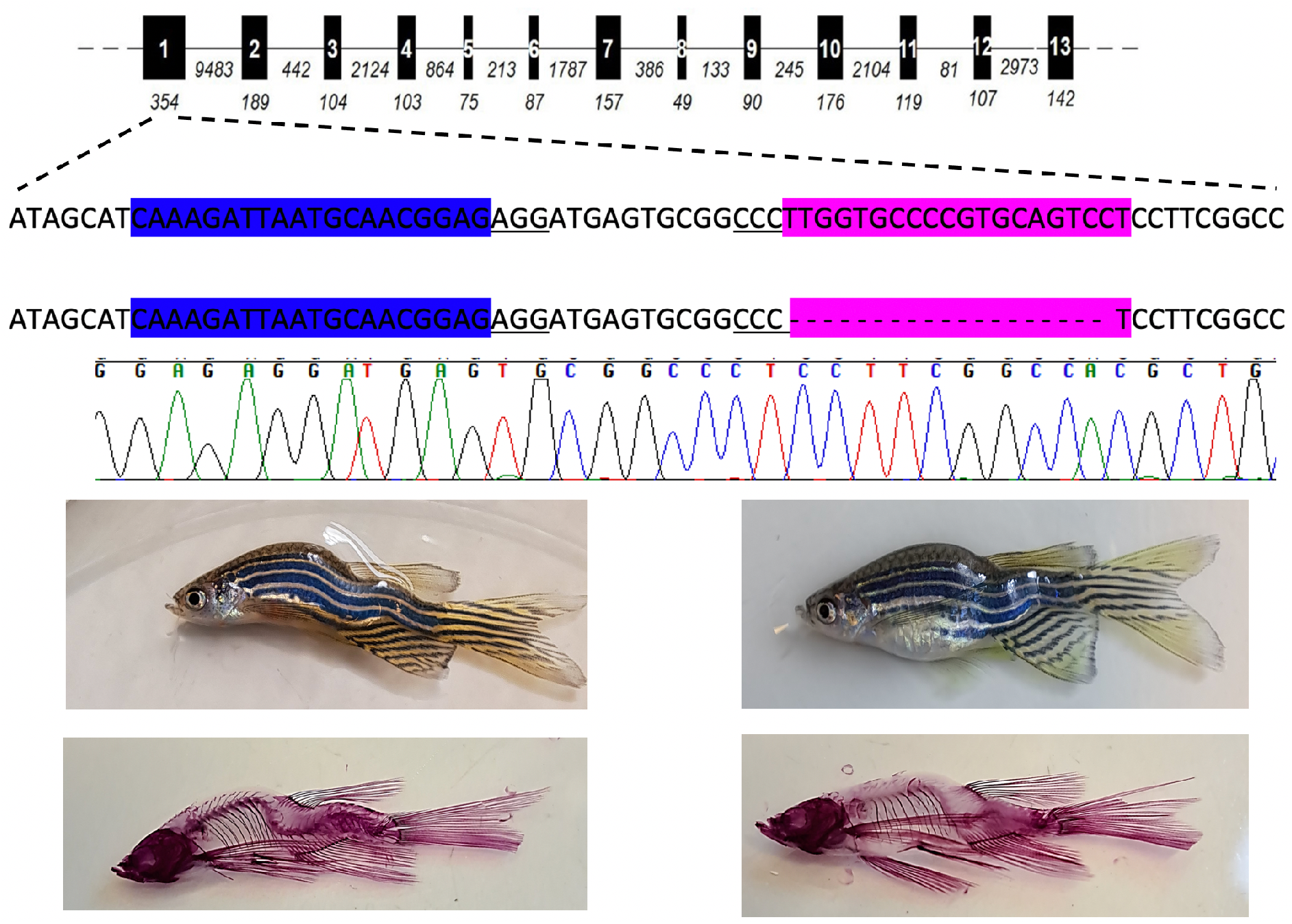
Scoliosis replicated in a mutant line encoding the mutation p.L82fsX24. Schematic of *ppp2r3b* showing the genic location of two sgRNAs used to target exon 1, highlighted in blue and pink, respectively. Deletion of 19bp is indicated by ‘-’ symbols and the confirmatory sequence chromatogram is shown. Representative examples of two 3 month old zebrafish (above) stained for Alizarin red (below).

### Kyphoscoliosis in *ppp2r3b* mutants is associated with reduced bone mineralisation of vertebrae

To investigate this phenotype further, we analysed bone mineralisation and cartilage formation in *ppp2r3b^−/−^* mutants. Alizarin red staining showed that the gross structure of all vertebrae was normal without the characteristic wedging of vertebrae that has been reported previously (Grimes et al. 2017), even at the sites of curvature (Figure 6A,B). Precaudal vertebrae 5-13 include a neural spine, which projects dorsally, and two ventrally located ribs, while the caudal vertebrae include neural and haemal spines which mirror one another in size. The ratio between the lengths of the neural and haemal spines within the caudal region showed that they were approximately equal in length in homozygous mutants as in wild-types and heterozygotes (Figure 6C). Within the precaudal region, the ribs are approximately 2.5 times longer that the neural spines (dorsal:ventral ratio of 0.4), however, we found that the ribs were relatively shorter in *ppp2r3b* homozygotes as compared to wild-types or heterozygotes (Figure 6C), suggesting a defect in patterning and/or ossification. We also noted a marked reduction in Alizarin red staining intensity throughout the vertebral body and spines/arches, which was uniform across all vertebrae in caudal and precaudal regions (Figure 6B).

To investigate bone formation in more detail, we performed microCT scanning to compare skeletal tissue parameters of vertebrae at 36 dpf which represents the onset of scoliosis. Remarkably, this analysis showed that multiple holes were apparent throughout the mutant vertebrae (Figure 8A). These represented sites of reduced bone mineral density (BMD) which was 32% lower in mutants as compared to wild-type, when quantified throughout the cortical bone (Figure 8B). Consistent with this, tissue mineral density was also significantly reduced. However, the overall dimensions of the vertebrae, including length and diameter, were not affected suggesting a specific effect on bone mineralisation rather than morphogenesis. qRT-PCR of RNA extracted from the trunk at 36 dpf did not reveal any changes in the expression of a panel of key bone cell markers (Table 1). Histological analyses of osteoclasts and osteoblasts did not reveal differences between wild-type and mutant (Figure 9)-Mallory’s trichrome staining showed similar numbers of vacuolated chordoblasts (osteoblasts) and squamous chordoblasts within the notochord centra and sheath, respectively. Furthermore, tartrate-resistant acid phosphatase (TRAP) staining demonstrated staining in neural and haemal arches, which was similar in wild-type and mutant, although no staining was detected within the vertebrae centra which is where the reduced BMD was observed previously. These data are subject to technical limitations relating to the proportionally small number of bone cells in whole zebrafish tissues at this stage and the non-quantitative nature of histological methods and are therefore not conclusively negative. In future, it will be necessary to analyse these cell types in more detail.

**Figure 8.**
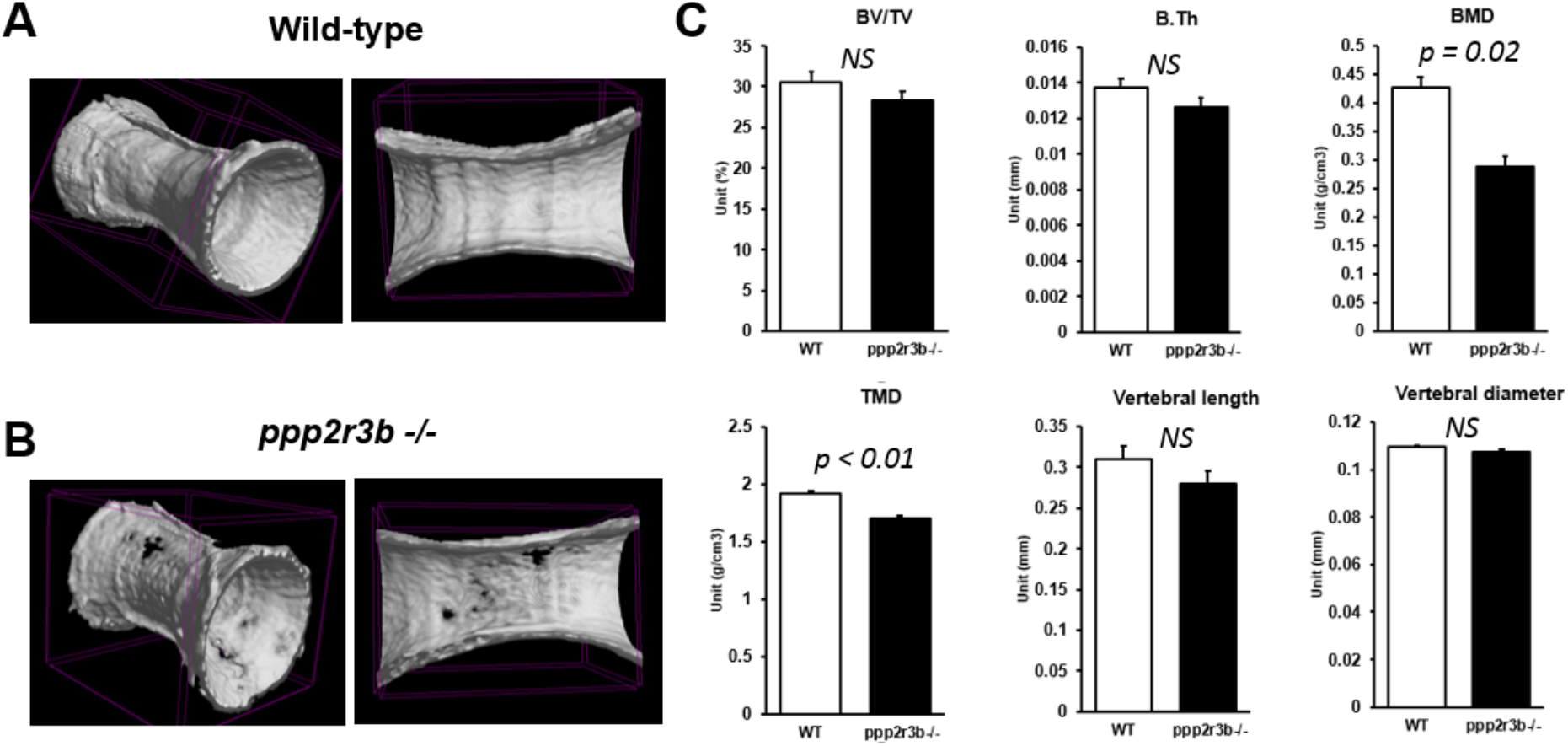
Reduced mineral density of vertebral cortical bone in *ppp2r3b^−/−^* mutants. (**A,B**) Representative 3D and cross-sectional images from microCT scans of vertebrae in wildtype and*ppp2r3b^−/−^* mutant zebrafish at 36 dpf. Note holes are apparent in the mutant vertebrae. (**C**) Quantification of a selection of cortical bone parameters measured in wild-type and *ppp2r3b^−/−^* mutant vertebrae (*n=3*) BV/TV, bone volume/tissue volume; B.Th, bone thickness; BMD, bone mineral density; TMD, tissue mineral density. *P-values* are given (*t-test*). NS, not significant. Average tail length for both mutant and wild-type animals was 12.6 mm with no difference between the two groups (*p-*value = 0.15, *t-test*). TLs of the two representative animals shown in **A** and **B** were 12.912 and 12.907 mm, respectively.

**Figure 9.**
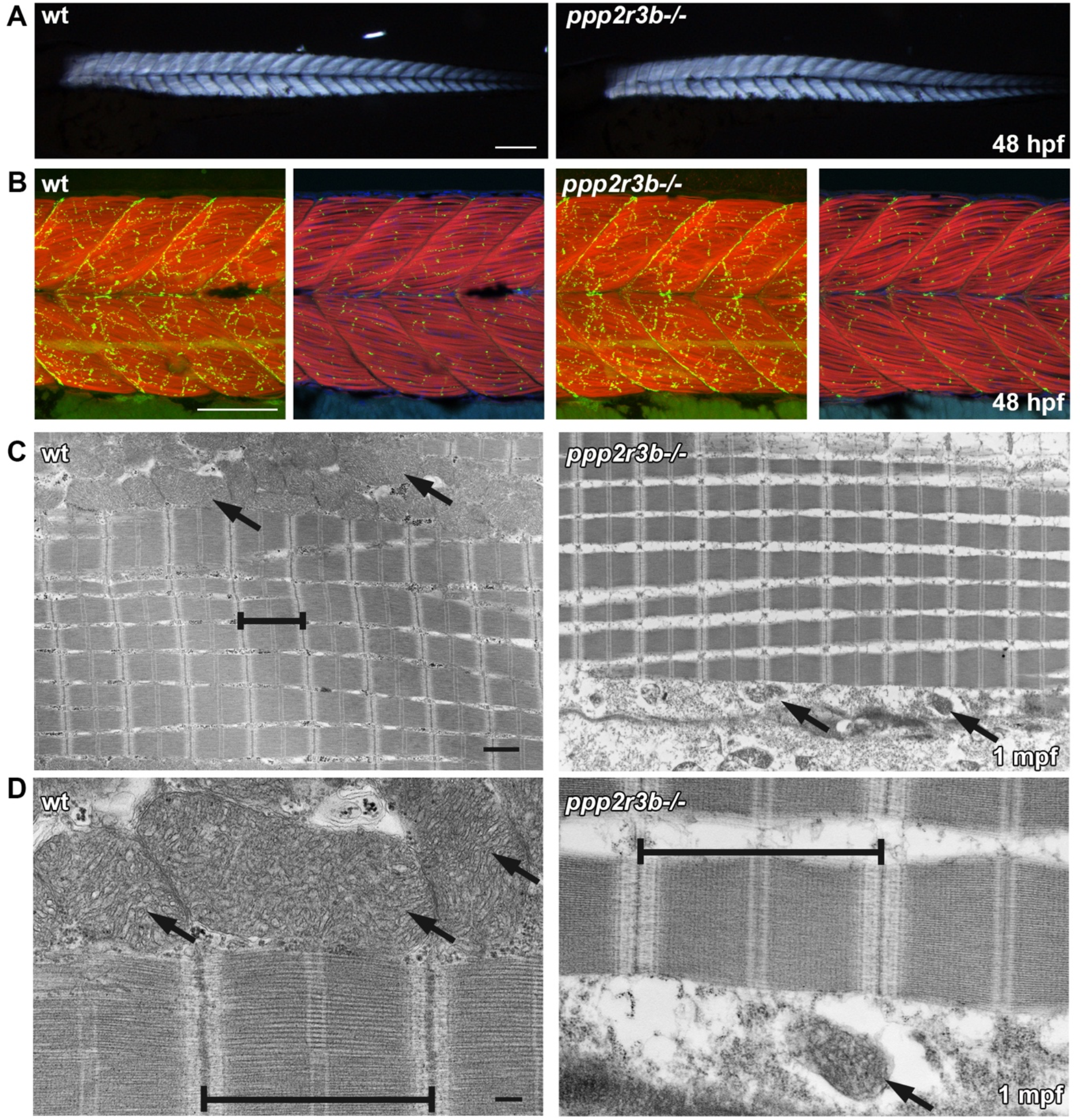
*ppp2r3b* mutants display abnormal mitochondria formation during adult stages of development, whilst muscle formation remains intact. (**A**) Skeletal muscle birefringence at 48hpf shows comparable muscle integrity between *ppp2r3b* mutants and wildtype siblings. Scale bar 200 um. (**B**) Muscle fibres and motor neuron synapses appear normal in mutants compared to wildtype siblings at 48hpf, analysed by staining for F-Actin (Red) and Acetylcholine receptors (AChR, green) respectively. Left panels show a z-stack projection of F-Actin with AChR, right panels show a single focal plane of F-Actin/AChR/DAPI. Scale bar 100 um. (**C**) Transmission electron micrographs indicate normal sarcomeric assembly (brackets) in juvenile mutants (1mpf) compared to wildtype controls, however mitochondria (arrows) are noticeably malformed and less abundant. Scale bar 1um. (**D**) High magnification electron micrographs showing detailed images of the sarcomeres (brackets) and mitochondria (arrows) in wildtype compared to mutant adolescent muscle samples. Scale bar 200 nm.

### Abnormal mitochondria associated with axial muscle in *ppp2r3b* mutants

Given the link between progressive spinal curvature and proprioception, and the expression of *PPP2R3B/ppp2r3b* that we detected in neurons, muscle and bone, we monitored muscle fibre and neuromuscular junction formation in our mutant fish. Initially we analysed skeletal muscle birefringence at 48hpf, an established indicator of muscle fibre integrity based on polarised light transmission through striated fibres. No obvious changes in birefringence signal were detected between *ppp2r3b* mutants and wildtype siblings (Fig. 9A). To challenge a role for PR70 in neuromuscular development, embryos were further stained for F-Actin and Acetycholine receptor (AChR), using fluorescently conjugated phalloidin and α-bungarotoxin. Muscle fibres and AChR localisation appeared organised and indistinguishable between *ppp2r3b^−/−^* embryos and wild-type siblings (Fig. 9B). Thus, PR70 is not required for normal neuromuscular development.

To evaluate muscle development during juvenile growth, at a point when scoliosis has manifested, transmission electron micrographs were produced from muscle biopsies. *ppp2r3b^−/−^* fish displayed normal myofibril organisation and sarcomeric assembly (Fig. 9C&D), and consistent with this, qRT-PCR analyses showed that expression of key muscle markers was unchanged (Table 1). However, a marked reduction in mitochondrial content was observed in *ppp2r3b* mutants (Fig. 9C). Closer examination of the mitochondria showed general dysmorphic character in the mutants with undefined cristae and smaller overall size (Fig. 9D). Taken together, these data suggest that *ppp2r3b*, whilst not required for the formation of muscle, is required to maintain muscle mitochondrial health.

## Discussion

In this study, we have created a targeted mutation in *ppp2r3b* in zebrafish which resulted in juvenile onset, progressive and fully penetrant kyphoscoliosis closely mirroring IS found in patents with TS, as well as abnormal mitochondria in the vicinity of axial musculature. Collectively the following pieces of information indicate that this results from a loss of PR70 protein function. While we did not see any differences in transcript levels in mutants by qRT-PCR and no specific antibody is available to detect PR70 protein, we did show that all detectable isoforms of *ppp2r3b* included exon 2, which carries the engineered frameshift mutation. This is located at residue 31 of the encoded protein which would therefore lead to deletion of two EF-hand motifs located within the C-terminal half of the protein which are essential for calcium binding and protein function (Dovega et al. 2014). Furthermore, protein sequence analysis does not indicate that there are any translation initiation codons downstream of the mutation which could produce protein product not involving the frameshift mutation. Another consideration relates to potential off-target mutations produced by the gene-editing process. Having bred the mutation out for a number of generations, any putative off-target mutations would be lost and certainly not be expected to segregate with the observed phenotype. By contrast, kyphoscoliosis segregated perfectly (100% penetrance) with the frameshift mutation over hundreds of animals and many generations, indicating linkage to the mutation. We note that we did also attempt to import both mutant alleles listed as available through the Zebrafish Mutation Project, but founders did not survive or breed possibly because of reduced embryo quality and/or passenger mutations.

The zebrafish has emerged as a powerful model to study IS (Grimes et al. 2016). By generating a zebrafish mutant in *ppp2r3b*, we have been able to investigate the requirement for this gene beyond larval stages, thereby revealing a requirement for normal bone mineralisation. Two key findings in relation to the potential cellular mechanism of pathogenesis were: 1) the identification of prominent *ppp2r3b* gene expression in somites and axial muscle in contrast to very limited expression within the vicinity of bone, and; 2) the identification of mitochondrial defects similar to those previously reported in muscular dystrophy models which followed normal muscle formation and occurred in the absence of muscle fibre (Z-disc) defects (Percival et al. 2013). Both the skeletal and muscle defects that we report were degenerative in nature. Collectively, these observations suggest that kyphoscoliosis occurs secondarily either to defects in proprioception driven by abnormal muscle function and/or because of disrupted communication between muscle and bone. The results from our electron microscopy studies suggests for the first time that defects in mitophagy are the primary cause of priopriocepion defects leading to loss of bone integrity. These are hypotheses that we will test in future.

At the molecular level, *PPP2R3B* encodes the DNA origin of replication complex (ORC) regulator, PR70 (Yan et al., 2000; Dovega et al. 2014; van Kempen et al. 2016.). Core ORC components, such as CDC6, are mutated in Meier-Gorlin syndrome which features a variety of skeletal defects including scoliosis (de Munnik et al. 2012; Bicknell et al. 2011). However, to our knowledge, nothing is known about the molecular or cellular function of the ORC in skeletogenesis. A zebrafish *cdc6* mutant has been reported, but post-larval phenotypes are potentially confounded by severe early embryonic defects and no skeletal features were reported (Yao et al. 2017). There was a recently identified role in autophagy as well (Pengo et al. 2017). The mouse homologue *Ppp2r3d* (mgi 1335093) appears to be an orthologue, in addition *Ppp2r3a* (mgi 2442104) is reported to have decreased bone mineral content. The zebrafish model that we report here provides a basis to investigate the molecular and cellular mechanisms of pathogenesis in future.

What is the potential relevance of this work to human disease? Turner syndrome (TS) is one of a group of relatively common aneuploidy disorders caused by partial or complete monosomy of the sex chromosomes (45X,O). For most genes on the X chromosome, only a single copy is required for normal development as they are subject to X-inactivation. A notable exception involves the pseudoautosomal region (PAR), which is common to both the X and Y chromosomes, and undergoes recombination and escapes X-inactivation. As such, it is likely that many features of TS are caused by haploinsufficiency for genes located within the PAR. TS is associated with a variety of musculoskeletal defects. This includes short stature, an abnormal jaw (micrognathia), as well as an increased prevalence of muscular dystrophy and scoliosis (Hanew et al. 2018; Ricotti et al. 2011; Day et al. 2007; Kim et al. 2001; Verma et al. 2017; Ferrier et al. 1963). The latter resembles IS and is progressive in nature with one study reporting a median onset of 9 years 11 months (Day et al. 2007). It is therefore possible that our findings in this work may be relevant to the pathogenesis of scoliosis in TS.

## Materials and methods

### *In situ* hybridisation

7μm paraffin sections were obtained from the Human Developmental Biology Resource (http://www.hdbr.org/). Riboprobes were synthesized with Digoxigenin-UTP RNA labeling kits (Roche) from amplified fragments of each gene using the following primers: *GFP* (553bp): EGFP_F, CGACGTAAACGGCCACAAG; EGFP_R, CTGGGTGCTCAGGTAGTGG, using pEGFP-N1 plasmid as template. *PPP2R3B* (556bp): PPP2R3B_F, CTTCTACGAGGAGCAGTGCC; PPP2R3B_R, TTTACACGAGCCGCGGTG. *SOX10* (561bp): SOX10_F, AGCCCAGGTGAAGACAGAGA; SOX10_R: TCTGTCCAGCCTGTTCTCCT. Template for PCR amplification of *PPP2R3B* and *SOX10* was human cDNA. The *SOX9* probe has been reported previously (Lai et al., 2003). Human embryonic samples were fixed in 10% (w/v) neutral-buffered formalin solution (Sigma-Aldrich) and embedded in paraffin wax before sectioning. ISH was performed in 300 mM NaCl, 5 mM EDTA, 20 mM Tris-HCl, 0.5 mg/mL yeast tRNA, 10% dextran sulfate, 1x Denhardt’s solution, and 50% formamide with digoxigenin-incorporated riboprobes at 68 C°. Posthybridization slides were incubated with anti-digoxigenin conjugated with alkaline phosphatase (Roche) diluted 1:1,000 in 2% newborn calf serum. Expression patterns were visualized with a Nitro-Blue Tetrazolium Chloride/5-Bromo-4-Chloro-3-Indolyphosphate p-Toluidine Salt (NBT/BCIP) system (Roche). Sections were mounted with Vectamount (Vector laboratories) and analyzed with a Zeiss Axioplan 2 imaging system.

### Zebrafish CRISPR/Cas9 mutagenesis using Cas9 plasmid

To create indels in the zebrafish *ppp2r3b* gene, the following sgRNA sequence was selected for targeting (5’ GGAATGCTTTCACTTAAGGC<u>TGG</u> 3’-PAM sequence is underlined). To create the sgRNA, the following 5’ phosphorylated oligonucleotides (5’ **TAGG**AATGCTTTCACTTAAGGC 3’ and 5’ **AAAC**GCCTTAAGTGAAAGCATT 3’) were denatured at 95°C and annealed by cooling to 25°C using a 5°C/min ramp. Oligos were subsequently cloned into pDR274. To generate sgRNAs, the plasmid was linearsied using *Dra I* and transcribed using the Megashortscript T7 kit (Invitrogen, AM1354). Capped RNA encoding Cas9 was synthesised from *XbaI-*linearised pT3TS-nCas9n plasmid using the T3 mMessage mMachine Kit (Ambion). The Poly(A) Tailing Kit (Ambion, AM1350) was used for polyA tailing of RNA. Both sgRNA and Cas9 RNA were purified using the RNeasy mini kit (Qiagen, 74104), which were subsequently co-injected (25 ng/μl sgRNA, 200 ng/μl Cas9 RNA) into zebrafish embryos at the 1 cell stage. To confirm successful mutagenesis total genomic DNA was extracted from individual 24 hpf embryos in 50mM NaOH at 95°C for >10 minutes, which was subsequently neutralised in 1 M Tris-HCl, pH 8.0. PCR was performed using the Phire Animal Tissue Direct PCR kit (Thermo Scientific F-140WH), which was subject to direct sequencing, T7 Endonuclease I assay (T7EI; NEB M0302L) or restriction enzyme digestion using *Mse I* (NEB R0525L). Splicing analysis of exon 2 was done using primers in exons 1 and 3 or exons 1 and 7, generating RT-PCR prodcuts of sizes 491 bp and 1000 bp, respectively. Primer sequences were as follows: ZFcDNA ppp2r3b Exon1_F aagggcacaagcactttgat, ZFcDNA ppp2r3b Exon3_R tttctccacattgccacaaa, cDNA_ppp2r3b Exon_1_F cttttggttgagtgagccgag and cDNA_ppp2r3b Exon_7_R atgccgcattaggtctctctg.

### Zebrafish CRISPR/Cas9 mutagenesis using RNPs

Three *ppp2r3b* targeting ribonucleotide protein (RNP) complexes were injected into zebrafish embryos to disrupt *ppp2r3b* function, increasing the likelihood of recovering targeted mutations. 2nmol crRNAs were synthesised against exon1 (IDT, CD.Cas9.SQVS6625.AL: CAAAGATTAATGCAACGGAG), (IDT, CD.Cas9.SQVS6625.AA: AGGACTGCACGGGGCACCAA), and exon 3 (IDT, CD.Cas9.SQVS6625.AI: ATGGAGCTTTCCAATAGAAT). RNP complexes were established following an adapted protocol from Kroll et al 2020. In brief, 2nmol crRNA for each target was resuspended in 10μl duplex buffer (IDT, #11-01-03-01). 5nmole tracrRNA (IDT,#1072532) was resuspended in 25μl Duplex buffer. Each crRNA was annealed to the tracrRNA by combining 1μl of crRNA with 1μl of tracrRNA and 1.5μl of duplex buffer, components were incubated at 95°C in a thermocycler. The Cas9 was then complexed to the annealed guide RNAs by adding 1μl of 10μg/μl Cas9 endonuclease (Alt-R S.p. HiFi Cas9 nuclease, IDT, #1081060) to each of the 3.5μl guide RNA solutions and incubated at 95°C for 5 mins. All three complexed RNPs were then pooled, mixed, aliquoted and stored at −20°C until required. 0.5nl of pooled RNPs were intracellularly microinjected into wildtype (AB x TupLF) embryos at the one-cell stage and embryos allowed to develop until adulthood.

### Morphometric analyses

Zebrafish were fixed in 10% neutral buffered formalin (NBF) for 24 h and stored in 70% ethanol until scanning. Alcian blue and Alizarin red staining was performed as described previously (Edsall et al. 2010). μCT analysis of cortical bone parameters was performed on the I st caudal vertebra (SkyScan 1172, Bruker, Belgium). A negative offset of around 10 sections (0.01 mm) was set in the selection of the vertebral bone. The μCT scanner was set at 40 Kv and 250 μA using no filter and a pixel size of 1.85 μm. Analysis of vertebral bone was performed ‘blind’. The images were reconstructed, analysed and visualised using SkyScan NRecon, CTAn and CTVox software, respectively. Bone mineral density (BMD) and tissue mineral density (TMD), were calibrated and calculated using hydroxyapatite phantoms with a known density. For all analyses presented in the manuscript, mutant and wild-type animals were matched to tail length.

### Analysis of muscle in zebrafish

Muscle birefringence in live embryos was visualized using a Nikon SMZ1270 dissecting microscope with a C-POL polarizing attachment. Embryos were immobilized in 3% methylcellulose and images acquired. Electron microscopy and staining of muscle and neuromuscular junctions was performed as described (Osborn et al. 2017).

### Ethics Statement

All animal procedures were authorised by Home Office Licence 70/7892 and work with human foetal tissues was authorised by the National Research Ethics Committee (REC reference, 08/H0712/34+5 and 18/LO/0822)

**Figure 10.**
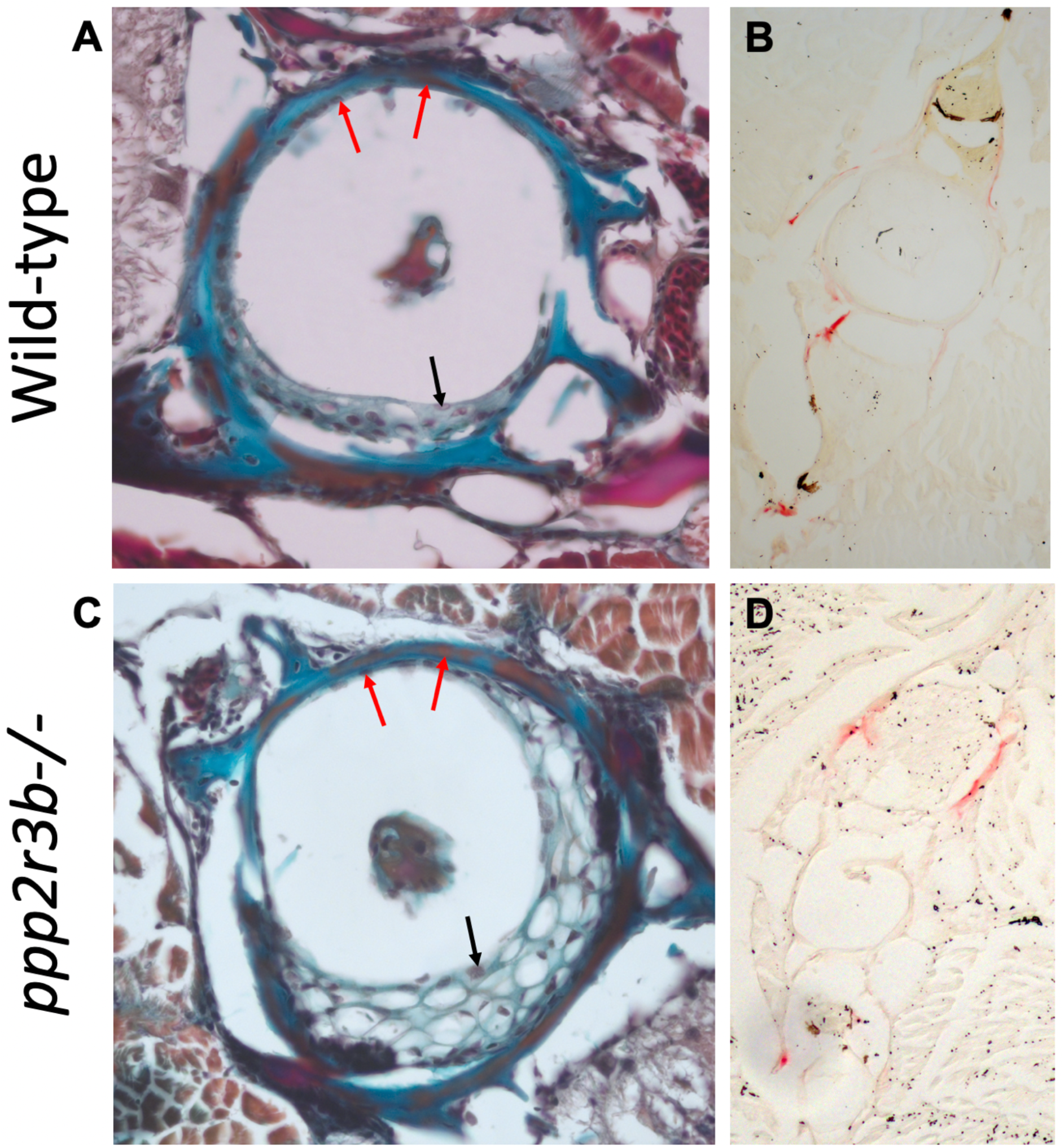
No obvious changes in osteoclast or osteoblast staining in vertebrae. (**A,C**) Mallory’s trichrome staining of vertebra centra. Black arrows show examples of vacuolated chordoblasts (osteoblasts) within the vertebra centrum while red arrows show squamous chordoblasts within the notochord sheath. (**B,D**) tartrate-resistant acid phosphatase (TRAP) staining in transverse sections from wild-type and *ppp2r3b−/−* mutant zebrafish at 36 dpf. Red signal indicates TRAP staining in neural and haemal arches, but no staining was detected within the vertebrae centra.

## Acknowledgements

This work was funded by a Medical Research Council New Investigator Research Grant (MR/L009978/1), Action Medical Research Project Grant (GN2595) and Wellcome Trust Collaborative Award in Science to DJ. This research was also supported by the National Institute for Health Research Biomedical Research Centre at Great Ormond Street Hospital for Children NHS Foundation Trust and University College London. The human embryonic and fetal material was provided by the Joint MRC/Wellcome Trust (MR/R006237/1) Human Developmental Biology Resource (www.hdbr.org).

## Notes

### Competing Interest Statement

The authors have declared no competing interest.

### Summary of Updates

We have modified the discussion and broad interpretation of disease relevance, and have included an additional zebrafish line with a separate mutation in ppp2r3b.

